# Developmental stage-specific responses to extreme climatic events and environmental variability in great tit nestlings

**DOI:** 10.1101/2025.10.06.680663

**Authors:** Devi Satarkar, David López-Idiáquez, Irem Sepil, Ben C. Sheldon

## Abstract

Climate change poses a pervasive threat to many aspects of natural systems, and while impacts of changes in average conditions have been extensively studied, the effects of increased climate variability, and extreme events, on natural populations are less understood due to the challenges of studying these rare occurrences. Using 60 years of life-history data from over 83,000 individuals, and historical daily climate records, we show that developmental stages in wild great tits (*Parus major*) differ in their sensitivity to extreme climatic events (ECEs). Exposure to extreme cold events during the first week of development is particularly detrimental to fledging mass, while extreme rain events have a stronger negative impact as chicks grow older and their energetic requirements increase. Synergistic effects of ECEs and average climatic conditions can be particularly severe, exacerbating the challenges faced by these birds. Our findings indicate that combined exposure to extreme heat and heavy rainfall during early development is associated with a predicted reduction in fledging mass by up to 27%. Additionally, birth timing may further modulate these effects, since late-season broods exposed to frequent hot ECEs during early development are predicted to fledge nestlings up to 4.27 standard deviations (35%) lighter than broods laid earlier in the season. Moreover, phenotypic plasticity has enabled many similar populations to shift towards an overall earlier laying date, which may have increased susceptibility to cold extremes during development. However, our analyses suggest that the benefits of being part of an early-laid clutch within a season may, to some extent, offset the negative effects of extreme climate on fledging mass and apparent survival. In climate scenarios where ECEs are predicted to increase in frequency, duration, and severity, these developmental stage-specific insights are crucial for understanding how climate change may be influencing wild avian populations.

## Introduction

Anthropogenic climate change has warmed the atmosphere at an unprecedented rate in recent decades, posing a profound and multifaceted threat to biodiversity (Urban et al., 2016). Animals are exhibiting diverse responses to changing average climates at the individual and population level, adjusting their phenology, demographic patterns, and spatial distributions (Johnston et al., 2019; Thackeray et al., 2016). For instance, there is well-documented evidence that plant and animal populations are shifting their ranges poleward or to higher altitudes in response to changing average climatic conditions (Lenoir & Svenning, 2015).

However, climate change manifests not only through shifts in mean conditions but also through increased climate variability, leading to changes in the frequency, intensity, and duration of extreme climatic events (ECEs) (Seneviratne et al., 2012). ECEs are deviations from typical climate patterns, often characterised by climatic conditions that fall in the tails of historical distributions (van de Pol et al., 2017). They present a unique challenge to ecological systems because species’ responses can be shaped by changes in both mean climate and patterns of ECEs, alongside their interactions (Lawson et al., 2015). Indeed, fluctuations in climate variability may have stronger effects on organisms than gradual shifts in average conditions, likely because they suddenly expose individuals to environments that they would have otherwise experienced and adapted to over larger timescales, and thus may exceed their physiological limits (Vasseur et al., 2014).

Global climate predictions indicate a virtually certain rise in the frequency and intensity of ECEs (IPCC, 2023), which is already evident worldwide, exemplified by more frequent and prolonged heat waves (Perkins-Kirkpatrick & Lewis, 2020) and intensified extreme precipitation events (Tabari, 2020). This escalating trend in ECEs necessitates an urgency to understand their ecological ramifications. However, the very nature of ECEs—their rarity and unpredictability—poses significant methodological challenges. Difficulties in eliminating confounding variables and achieving statistically robust assessments have led to a predominant focus on short-term impacts, leaving the effects of multiple extreme events and long-term responses less understood (Bailey & van de Pol, 2016; Regan & Sheldon, 2023).

Despite these challenges, a few studies on animal populations have revealed how ECEs can disrupt life-histories and drive evolutionary change. Blue tits (*Cyanistes caeruleus*) experiencing extremely hot days while rearing chicks had fewer fledglings and increased selection for earlier breeding (Marrot et al., 2017). Reduced reproductive output was also observed in great tits (*Parus major*) that were exposed to multiple ECEs (Regan & Sheldon, 2023). Furthermore, survival in red-winged fairy-wrens (*Malurus elegans*) and white-browed scrub wrens (*Sericornis frontalis*) were more strongly linked to temperature extremes than average conditions, with carry-over effects of climate in prior seasons mediating size-dependent mortality (Gardner et al., 2017). In superb fairy-wrens (*Malurus cyaneus*), weather variables exerted counteracting effects on offspring body size across different timescales, illustrating the complexity of disentangling short-term and cumulative climatic impacts (Kruuk et al., 2015). Yet, most existing research has focussed on the isolated effects of ECEs. There is a pressing need for comprehensive studies that examine how multiple ECEs interact with other ecological factors and long-term climatic trends to influence wild populations.

Body mass at fledging has frequently been used as a proxy for fitness prospects in nestling birds. Apart from integrating physiological condition, environmental constraints, and parental investment into a single metric, it is also frequently predictive of survival and reproductive traits (Both et al., 1999; Bouwhuis et al., 2015; Garant et al., 2004; Monrós et al., 2002; Perrins, 1965). Temperature fluctuations can significantly influence nestling growth rates, with both positive (Marques-Santos & Dingemanse, 2020; Matthysen et al., 2011; Mccarty & Winkler, 1999) and negative (Mainwaring & Hartley, 2016) effects observed depending on the context and species. Warmer temperatures may enhance growth by increasing food availability and reducing thermoregulatory costs, but extreme heat can lead to dehydration and reduced parental foraging efficiency. Observational studies of great tits have also shown that increased precipitation during spring leads to lower fledging weight, potentially due to reduced parental foraging effort during rainfall (Keller & Van Noordwijk, 1994; Radford et al., 2001). Experimental manipulations of nest microclimates have further demonstrated that higher nest temperatures can both enhance (Dawson et al., 2005) and hinder (Andreasson et al., 2018; Rodríguez & Barba, 2016b; Woodruff et al., 2025) growth rates in many cavity-nesting birds. These seemingly contradictory findings highlight how context-dependent temperature effects can be. Most of these studies have explored the impacts of average climate, with notable temperature effects during specific developmental windows. Moreover, there is large variation in local climatic conditions among study sites. Temperature rises may exceed species’ thermal limits and be more detrimental to growth and survival in warmer regions as compared to cooler habitats.

Given the projected increase in ECEs, understanding their effects on body mass is critical for predicting population-level responses to climate change. Our study system of great tits in Wytham Woods, Oxfordshire, UK, established in 1947 (Lack, 1964), is an appropriate site to examine ecological responses to environmental changes, including climate shifts and rare events, over a timeframe that spans significant variations in ecosystem dynamics. Great tits are passerines that inhabit temperate forests and breed during spring when they rear large broods on an insect diet (Lack, 1964). The rearing period of these altricial birds is particularly sensitive to temperature fluctuations, with an ectothermic phase in the initial few days, before they eventually attain thermoregulatory mechanisms (Rodríguez & Barba, 2016a). They exhibit rapid growth over a well-defined period, with nestlings growing to 10 times their body weight, from hatching to fledging, in approximately 21 days. This allows for precise targeting of specific developmental phases to understand how variable climates may influence particular stages of nestling growth.

Our long-term dataset, spanning more than 83,000 individual-level observations across 60 years analysed here, enabled us to detect rare ECEs and analyse their impacts on a natural population over ecologically meaningful time-scales—an important feature given the rarity and unpredictability of such events. We leveraged this to understand how exposure to ECEs during critical developmental periods may impact both short-term and long-term outcomes in this population. To this end, our study had three objectives. First, we aimed to assess the direct effects of the ambient climate, and ECEs on fledging weight, in two different developmental stages, hatchling (0-7 day old) and nestling (8-15 day old), which are expected to exhibit differential sensitivities to temperature fluctuations. Second, recognising that climatic factors rarely act in isolation, we explored how ECEs may interact with ambient climate and relative laying date, that is how early or late a clutch is laid within each breeding season compared to the seasonal average. These interactions are important for understanding how the impacts of ECEs may be influenced by concurrent variations in temperature, rainfall, or other environmental conditions. Earlier broods generally have access to more abundant resources (Charmantier et al., 2008; Verboven & Visser, 1998) and may better withstand environmental stresses, although this relationship is influenced by variation in both food timing and parental capacity. By examining variation in relative lay date and its interaction with exposure to extreme climatic events (ECEs), we assessed how phenological timing among broods within a season may modulate offspring responses to climate variability, which is crucial given that Wytham’s great tits have shifted towards earlier laying over the years (Charmantier et al., 2008). Finally, we investigated whether exposure to ECEs during development has an effect on apparent survival to recruit to the breeding population, to explore long-term consequences of extreme climate variability on population dynamics.

## Methods

### Study system

The great tit population in Wytham Woods, Oxfordshire, UK, has been systematically monitored since 1947 (Perrins, 1965). In this mixed-deciduous woodland, over 1000 artificial nestboxes have been provisioned for cavity-nesting passerines since the 1960s. These nest-boxes are visited at least once a week during the breeding season (April-June) to collect information on egg-laying date (date when the first egg is laid), hatching date, clutch size and number of fledglings. As part of our standard protocol, nestlings are fitted with unique metal rings from British Trust for Ornithology and weighed to the nearest 0.1 g at 15 days old, when their weight typically plateaus, providing a reliable measure of fledging mass (Bouwhuis et al., 2015). Parents are captured at the nest when nestlings are between 12 and 14 days, during the provisioning stage, and are individually marked if they have not been previously tagged. For this study, we analysed sixty years’ data from 1965 to 2024, using fledging mass and apparent survival (assessed as recruitment to breeding population in subsequent years) as variables of interest. Consistent with previous work on this system, we decided to use data from 1965 onwards to account for the population stabilising following the installation of new nestboxes in 1961 (Regan & Sheldon, 2023).

### Characterising ECEs during developmental periods

To examine the stage-specific impacts of climate on chick development, we focused on two distinct periods prior to the fledging weight measurement at 15 days old – the hatchling stage (0-7 days post-hatch) and the nestling stage (8-15 days post-hatch). We chose these periods due to their differential environmental sensitivities and physiological characteristics (Marrot et al., 2017). During early development, hatchlings are particularly vulnerable to temperature fluctuations due to their lack of feathers and limited thermoregulatory capacity. On the other hand, the nestling stage is characterised by improved thermoregulation but substantially higher food requirements.

We used daily temperature and precipitation data from the Met Office Hadley Centre datasets for central England (https://www.metoffice.gov.uk/hadobs/) and calculated the average temperature and rainfall during both the periods. We also used these data to define ECEs as events falling within the extreme 5% tails of the temperature and rainfall distributions observed over the study period (1965-2024). Specifically, we calculated daily deviations from the monthly means and used the 5th and 95th percentiles to establish cut-offs (Marrot et al., 2017; Regan & Sheldon, 2023). Hot ECEs were defined as days with temperature ≥+4.52°C above the monthly mean, cold ECEs as ≤ -4.49°C below the monthly mean, and rain ECEs as total rainfall in 24 hours ≥ 6.20 mm above the monthly mean. It is important to note that our definition allowed for multiple ECEs to be recorded on consecutive days if the extreme conditions persisted, as each day meeting the criteria was counted as a separate event. Using these thresholds, we quantified the frequency of ECEs during both the hatchling and nestling stages for each individual chick.

In order to explore whether the magnitude of the deviation in climatic conditions from normal is relevant, we also quantified “more extreme” ECEs using the 1% tails of the temperature and rainfall distributions, representing conditions much further from the long-term mean than the standard 5% definition. Consequently, the much higher thresholds for 1% hot, cold, and rain ECEs were daily temperature ≥ +6.27°C, ≤ –6.27°C, and rainfall ≥ 13.41 mm above the respective monthly means (values calculated for the entire 1965–2024 period).

### Statistical analysis

All analyses were conducted using R version 4.3.3 and linear and generalised linear mixed models were fitted using *lme4* (version 1.1.35.3). We conducted three sets of analyses: (1) Effects of ambient climate and ECEs on fledging mass, (2) interactions of ECEs with ambient climate and birth timing, (3) effects of ECEs on apparent survival. We examined data and model residuals for normality using histograms and Q-Q plots. Gaussian distribution was assumed for most models unless specified otherwise. We also checked for multicollinearity among model variables, confirming that variance inflation factors remained below 3 (Fox & Weisberg, 2018) (using the *performance* package (version 0.12.2)). Plots were constructed with predicted trends based on model estimates using the *ggeffects* package (version 1.5.2). All model outputs with details of predictors, estimates (β), standard errors, test statistics, and confidence intervals, are provided in the supplementary information.

### Effects of ambient climate and ECEs on fledging mass

Using 60 years of continuous life-history data, we analysed the fledging weight of 83,935 individual chicks (median (IQR); 18.5 g (17.6-19.4 g)). For each of the 11,609 broods, we matched average daily temperature and rainfall values calculated specifically for the two developmental stages (hatchling and nestling) according to the recorded hatchdate, which ensured that the climate predictors accurately reflected the environmental conditions experienced by each chick during specific developmental windows. We constructed individual-level linear mixed models for both the developmental periods in which fledging mass was the response variable and the average temperature or rainfall during the relevant period was the main predictor. All models included clutch size and laying date as covariates to account for their well-established effects on chick body mass. Non-independence of offspring raised in the same brood, and of broods raised in the same year, and at the same location, was taken into account by fitting random effects of year of birth, brood identity, mother identity and natal nest box.

For temperature, we suspected a nonlinear relationship with fledging mass based on exploratory plots and biological reasoning, since many physiological and ecological processes exhibit responses to temperature that involve thresholds, optima, or plateaus, and may not be well captured by strictly linear models. We used natural cubic splines with 5 degrees of freedom (via the ns() function in R’s *splines* package (version 4.3.3)) to flexibly model this effect. This approach helps capture curvilinear responses by fitting piecewise cubic polynomials that are joined smoothly at knot points. The choice of 5 degrees of freedom offered a balance between sufficient flexibility and model parsimony, to visualise plausible patterns between average temperature experienced during development and the subsequent body mass at fledging. For a detailed explanation of this methodology, see Harrell, 2015. For average rainfall models, a linear model provided a robust fit across the data range. While spline models offered flexibility here, they introduced instability in the fit where data was sparse at higher values.

We fitted separate models for exploring the effects of ECEs during the hatchling or nestling stage, by incorporating the number of ECEs as a fixed effect, along with average temperature, clutch size and laying date as covariates. Each type of ECE was incorporated in separate models due to collinearity between the number of cold and hot ECEs. We also constructed a spline model for hot ECEs as it had a better fit than a linear model. Random effects here were the same as before. All fixed effects were scaled to a mean of zero and standard deviation of one to allow for a direct comparison between effect sizes.

Additionally, to test for the effects of even more extreme ECEs, we constructed mixed-effects models within the same framework as described above, using binary indicators for the presence of at least one very extreme event (1% tail) and at least one extreme event (5% tail) during each developmental stage. These stage- and event-type–specific binary variables were derived from the same temperature- and rainfall-based definitions of ECEs, but coded as presence/absence rather than counts. As before, models were fitted separately for hot, cold, and rain ECEs at each stage using the same fixed covariates (average temperature for the relevant stage, laying date, clutch size) and random intercepts (birth year, brood identity, mother identity, natal nest box).

### Interactions of ECEs with ambient climate and birth timing

To examine how ambient climatic conditions influence nestling mass within the context of extreme climatic events (ECEs), we expanded our modelling framework by introducing interaction terms between climate variables and ECE metrics. Specifically, we tested whether the effects of average temperature vary with the presence and severity of rain ECEs, and whether the influence of average rainfall changes in the presence of hot or cold extremes. This was designed to reflect the ecological reality that climate variables seldom operate independently, and that the biological impacts of extreme climate can be shaped by prevailing temperature and precipitation patterns.

Furthermore, to assess the interplay between ECEs and resource availability, we incorporated relative lay date, calculated as the difference between individual lay date and the population average in the same year, as an interaction term with the number of ECEs. We acknowledge that resource availability is a complex process influenced not only by laying date, but also factors like prey phenology, weather, and habitat quality, which are challenging to quantify directly at this scale. Nevertheless, we used relative lay date as a proxy for seasonal shifts in resource dynamics, because earlier laying generally aligns with peak food abundance and can potentially afford parents greater flexibility in matching offspring development to resource peaks. This approach would allow us to investigate whether the timing of breeding within the season changes the degree to which nestlings are affected by extreme climatic variability. All interaction models used ECE variables with the 5% threshold.

### Effects of ECEs on apparent survival

Apparent survival for each individual was determined by assessing the local recruitment of the fledgling to the breeding population in subsequent years. Local recruitment was thus a binary variable with 1 for recruited and 0 if not known. This measure underestimates actual survival, because birds do emigrate from Wytham, or may breed locally without being captured, but emigration has been shown to be independent of fledging mass and unlikely to bias results (Bouwhuis et al., 2015). We analysed 83,935 individuals hatched between 1965 to 2024, with the latest recruitment to the 2025 breeding season.

To investigate the long-term effects of ECEs during the hatchling and nestling stages, we constructed generalised linear mixed models (GLMM) with local recruitment as the response and the number of ECEs (5% threshold) as the fixed effect. As before, average temperature and clutch size were incorporated as additional fixed effects, with birth year, brood identity, mother identity, and nest box as random effects. The GLMMs assumed a binomial distribution. We also constructed additional models where the laying date was accounted for as fixed effect.

For these models, we enhanced the predictive power to examine ECE effects by combining higher ECE frequencies into a single category, treating the number of ECEs as a categorical variable rather than a continuous one (Supplementary Table 7). This accounted for the rarity of individuals experiencing very high ECE frequencies and allowed us to detect significant effects of ECEs beyond a certain threshold (Regan & Sheldon, 2023).

## Results

### Effects of ambient climate and ECEs on fledging mass

Linear mixed models using natural cubic splines revealed significant nonlinear relationships between average ambient temperature (in both developmental phases) and fledging weight (Fig. 1A, 1B). In the hatchling phase, multiple spline terms were significant (e.g., β_1_ = 0.82 ± 0.15, t = 5.33, p < 0.001; β_3_ = 0.79 ± 0.13, t = 6.26, p < 0.001), indicating a flexible curvilinear relationship. A similar nonlinear pattern was observed during the nestling stage, where at least one spline term strongly predicted fledging mass (e.g., β_3_ = 0.84 ± 0.15, t = 5.42, p < 0.001). High average rainfall during both phases was associated with reduced fledging weight, with the effect being more pronounced in the nestling phase (hatchling: β = -0.063 ± 0.018, t = -3.467, p < 0.001; nestling: β =-0.143 ± 0.015, t = -9.109, p < 0.001; Fig. 1C, 1D).

**Figure 1:**
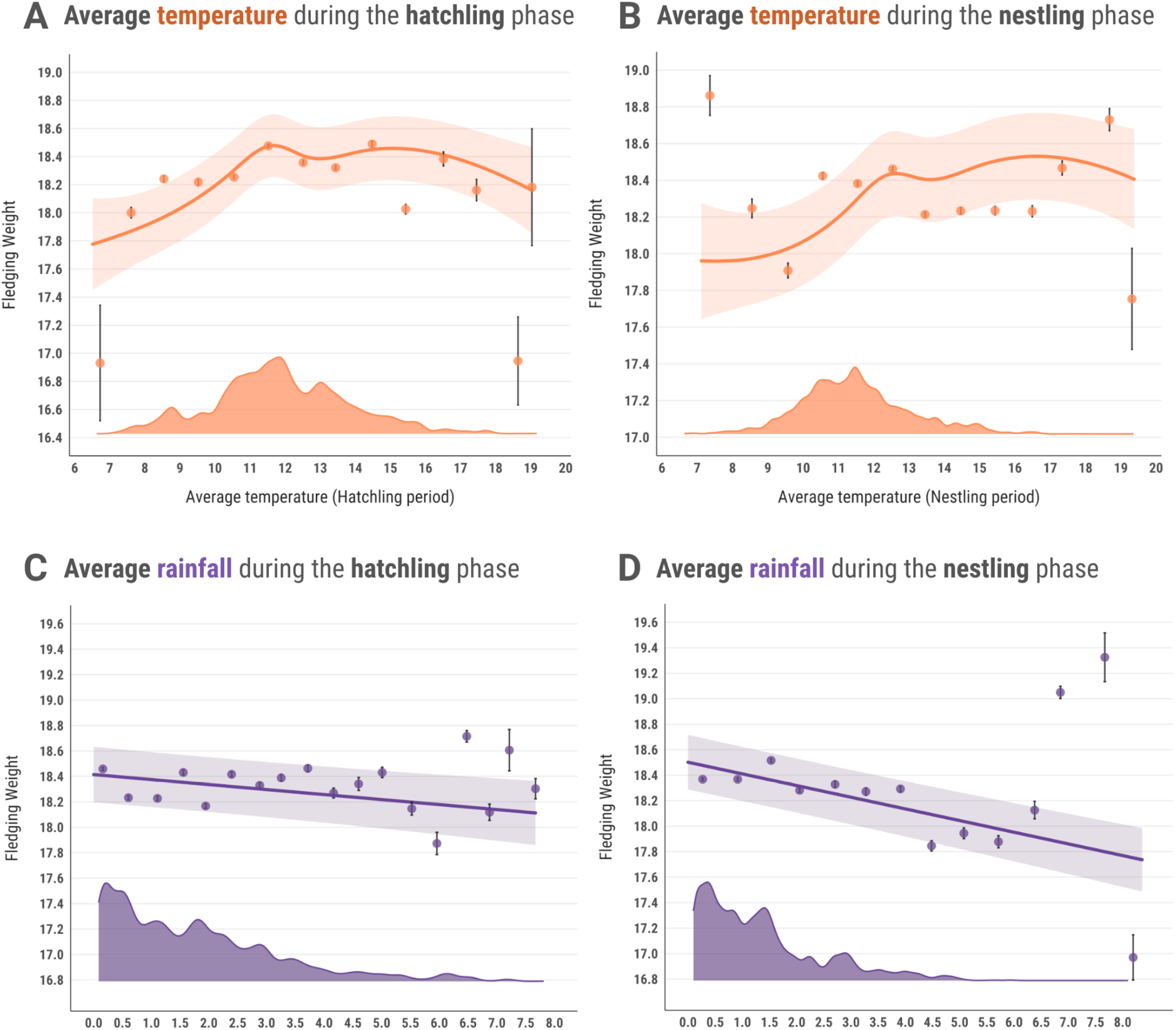
Associations between average temperature (°C) and fledging weight (g) in the hatchling (A) and nestling (B) stages, and between average rainfall (mm) and fledging weight (g) in the hatchling (C) and nestling (D) stages. Solid lines and associated ribbons indicate predicted trends and 95% confidence intervals from the models. The coloured dots and lines represent the average fledging weight ± standard error from observed data grouped in 1°C (temperature in orange) and 0.5 mm (rainfall in purple) bins respectively. The density plots above the x axes show the distribution of data points from observed data (*N* = 83,936 individuals).

Hot ECEs during the hatchling stage had no significant effect on fledging mass (β = –0.022 ± 0.023, t = –0.99, p = 0.32). In contrast, fledging mass increased nonlinearly with increased frequency of hot extreme events during the nestling stage, as indicated by a natural spline model (Fig. 2A). Predicted fledging weight remained relatively stable up to three hot days before increasing more sharply with further exposure. Chicks experiencing seven hot ECEs were predicted to weigh approximately 0.5 standard deviations (4.5%) more than those with no exposure.

**Figure 2:**
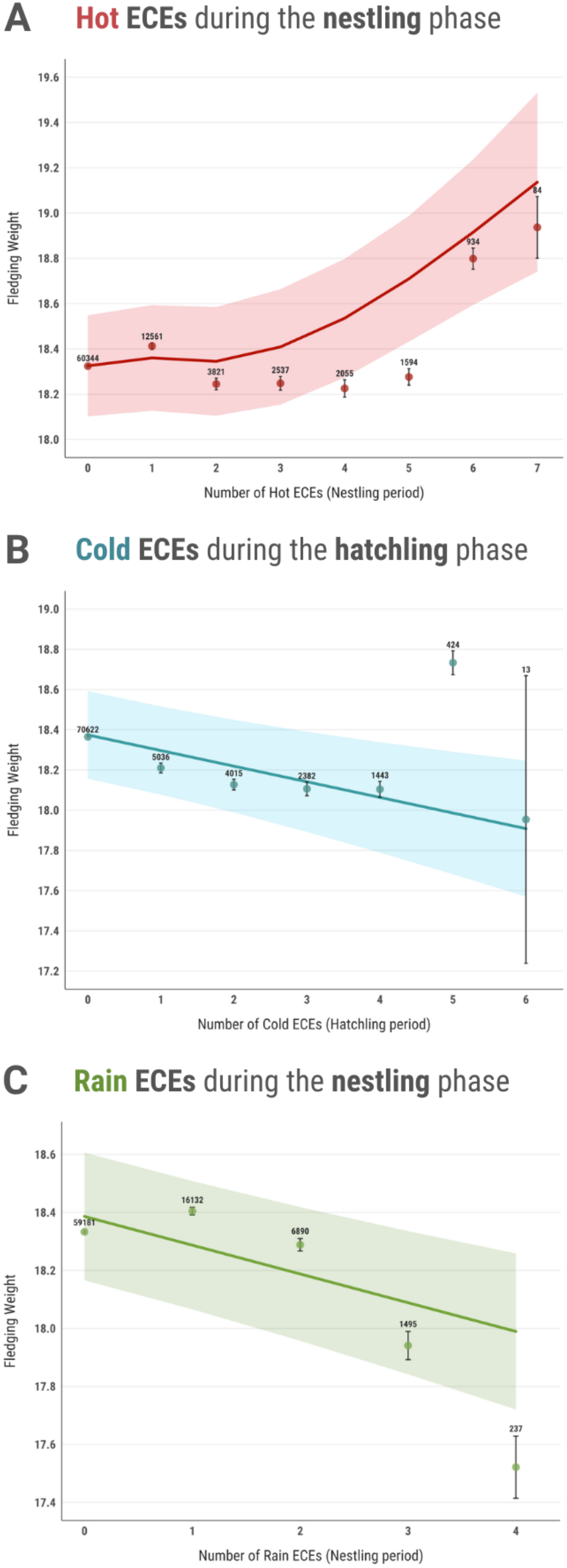
Predicted effects of (A) hot ECEs in the nestling phase, (B) cold ECEs in the hatchling phase, and (C) rain ECEs in the nestling phase on fledging weight (g). Solid lines and associated ribbons indicate predicted trends and 95% confidence intervals from the models. The coloured dots and lines represent the average fledging weight ± standard error for each frequency of ECE from observed data (*N* = 83,936 individuals). Predicted trends account for the model structure, incorporating fixed and random effects, which may cause predicted values to differ from raw summary statistics. Sample sizes for each frequency of ECE are indicated above the corresponding data points.

However, frequent cold ECEs during the hatchling period were associated with a significant reduction in fledging mass (Fig. 2B). The linear model predicted that chicks experiencing cold ECEs throughout the entire hatchling period (i.e., 6 consecutive days) would fledge at weights that were on average 0.28 standard deviations (SD) lower than those of chicks without cold ECE exposure. No significant effect of rain ECEs was observed during the hatchling stage (p = 0.14), but the nestling stage showed vulnerability to extreme precipitation (β = -0.073 ± 0.016, t = -4.536, p < 0.001), with a predicted average reduction of 0.4g (0.24 SD) in fledging mass of chicks that experienced 4 days of extreme rainfall compared to those with no rain ECEs (Fig. 2C).

Additional models using binary indicators for the presence of at least one very extreme event (1% threshold) yielded patterns of association broadly consistent with the frequency-based models above albeit with mostly non-significant and weaker effect sizes across developmental stages, especially for hot and cold ECEs. Very extreme rainfall across both hatchling (β = –0.163 ± 0.063, p = 0.010) and nestling (β = –0.149 ± 0.06, p = 0.013) stages was associated with reduced fledging masses, although these effects were somewhat weaker than those observed for the 5% rain ECEs. Full details of all count-based 5% ECE and binary 1% ECE model results are provided in Supplementary Tables 2 and 3.

### Interactions of ECEs with ambient climate and birth timing

Our models revealed significant interactive effects between the frequency of ECEs and ambient environmental conditions on chick growth outcomes, but only during the hatchling stage. While experiencing rain and hot ECEs during this early phase of development did not lead to any significant direct effects on fledging mass, their combined effects with ambient climate variables were pronounced. In particular, the detrimental impact of increased precipitation during this stage was amplified with increasingly frequent hot ECEs (hot ECEs × mean rainfall: β = -0.162 ± 0.022, p < 0.001; Fig. 3A). For example, at the highest frequency of hot ECEs (6 events), predicted fledging mass decreased by approximately 27%, corresponding to a 3.15 SD reduction as average rainfall increased from 0 to 8.36 mm. In contrast, chicks with no exposure to hot ECEs showed a decrease of only 0.08 standard deviations (0.7%) (Fig. 3A).

**Figure 3:**
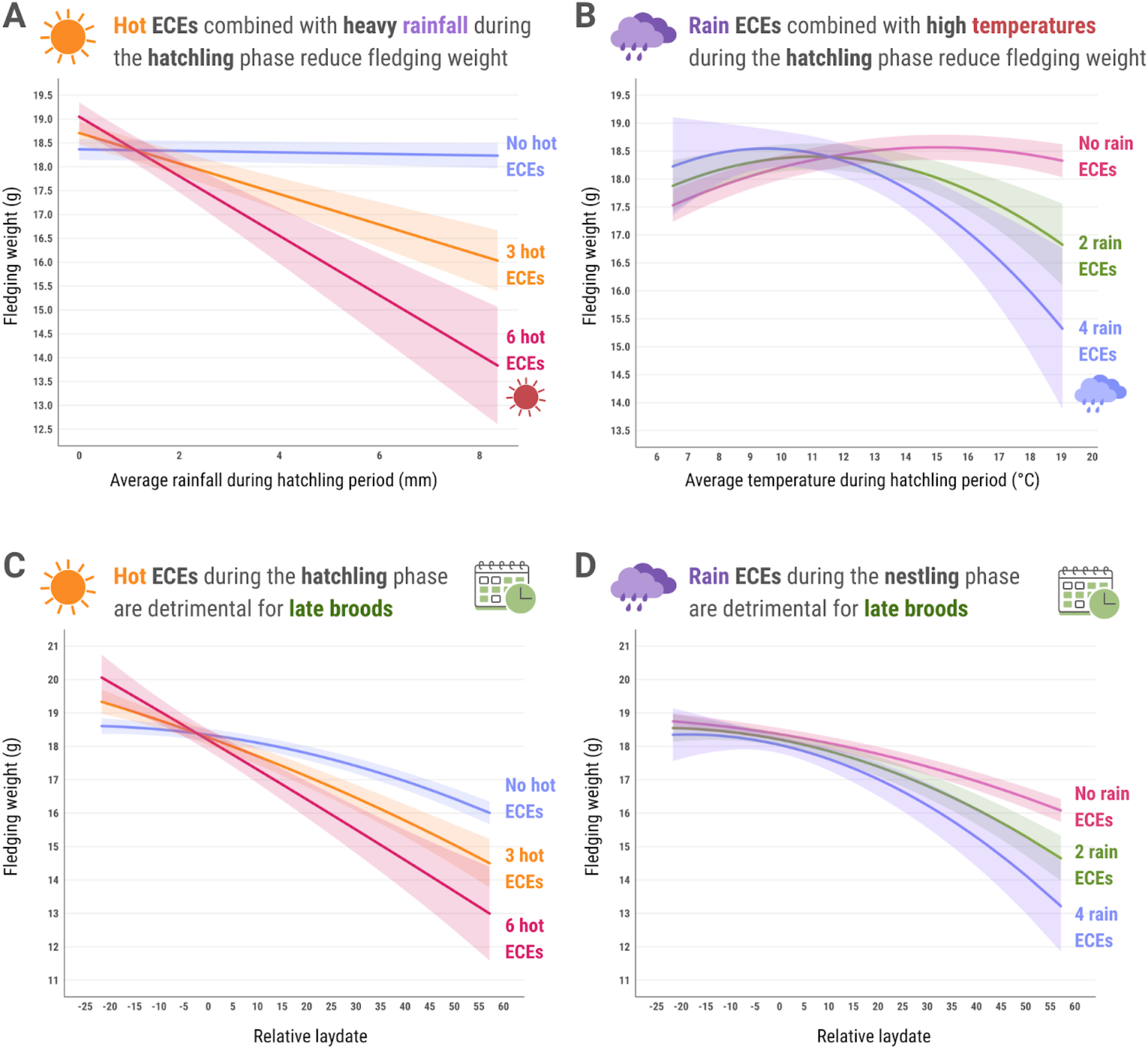
Predicted trends from interactions between (A) number of hot ECEs x average rainfall (mm) during the hatchling stage, (B) number of rain ECEs x average temperature (°C) during the hatchling stage, (C) number of hot ECEs during the hatchling stage and relative lay date, (D) number of rain ECEs during the nestling stage and relative lay date; relative lay date is the difference between individual lay-date and average lay date of the population in that year, negative values indicate relatively earlier clutches. Solid lines and associated ribbons indicate predicted trends and 95% confidence intervals from the models.

Furthermore, extreme precipitation during this stage narrowed the optimal temperature range for early development, exacerbating the negative effects of high temperatures when co-occurring (rain ECEs × mean temperature: β = -0.085 ± 0.017, t = -4.973, p < 0.001; Fig. 3B). Increasing rain ECEs from 0 to 4 at high average temperatures (∼19°C) led to about a 16% reduction in predicted chick mass (- 1.817 SD). These results highlight how combined stressors can synergistically influence thermoregulatory mechanisms in hatchlings.

Stage-specific interactions between extreme climatic events (ECEs) and environmental conditions were further modulated by hatch timing relative to the population’s seasonal average. Predicted fledging weights from our models revealed that extreme heat exposure during the hatchling stage may disproportionately impact relatively later broods (Fig. 3C). Estimates for the earliest broods (∼22 days earlier than the seasonal average) predicted a moderate increase in fledging weight of about 7.8% (+0.877 SD) from chicks that have not experienced any hot ECEs, to those that have experienced them throughout the hatchling stage. In stark contrast, chicks from later broods (∼57 days after the seasonal mean) showed a predicted decline of nearly 1.83 standard deviations (18.9%) under the same heat exposure. A predicted difference of approximately 4.27 SD (>35%) in fledging mass was observed between early and late broods exposed to six consecutive days of extreme heat (hot ECEs × relative lay date: β = -0.067 ± 0.012, t = - 5.181, p < 0.001; Fig 3C).

Similarly, rain ECEs during the nestling phase had a greater negative impact on broods born later in the breeding season. This interaction widened the fledging mass gap between early and late clutches. In the absence of rain ECEs, the predicted fledging mass difference between earliest and latest clutches was 1.61 SD (14.2%), but this disparity nearly doubled to 3.1 SD (27.9%) if nestlings experienced 4 rain ECEs (Fig. 3D). The accelerating decline in late broods with increasing rain ECEs is reflected in a significant quadratic interaction term (β = -0.014 ± 0.006, t = -2.334, p = 0.01).

### Effects of ECEs on apparent survival

We found evidence that recruitment probability (a proxy for survival) declined with increased exposure to high frequencies of extreme cold and rain. Data from 83,935 individuals (1965–2024) revealed an average recruitment probability of 9.1% over the study period. Exposure to cold ECEs during the hatchling stage reduced survival probability, with a significant negative effect observed for individuals experiencing a single extremely cold day (β = -0.241 ± 0.07, *z* = -3.273, *p* = 0.001). Survival probability was predicted to decrease by over 25%, from 8.0% for individuals with no cold ECE exposure to 5.8% for those exposed to four or more cold ECEs (β = -0.30 ± 0.128, z = -2.368, p < 0.05, Fig. 4C).

**Figure 4:**
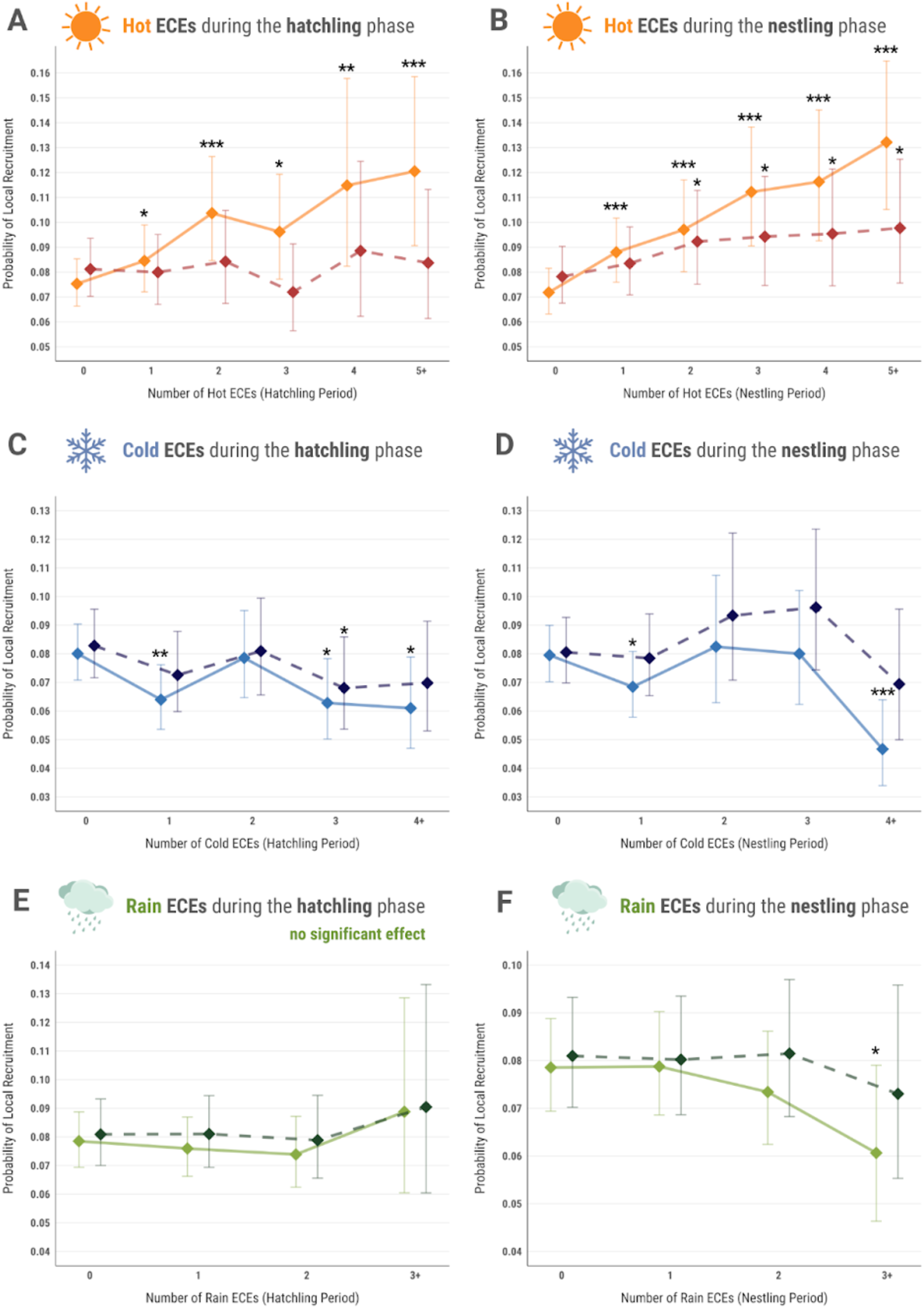
Associations between apparent survival probability (0 to 1) and (A) number of hot ECEs during the hatchling stage; (B) number of hot ECEs during the nestling stage; (C) number of cold ECEs during the hatchling stage; (D) number of cold ECEs during the nestling stage; (E) number of rain ECEs during the hatchling stage; (F) number of rain ECEs during the nestling stage. Shown are model predictions and associated credible intervals; *N* = 82,229 individuals. Solid lines represent predicted trends before laying date was accounted for in the model, whilst dashed lines correspond to model predictions when individual laying dates were included as fixed effect. Significant effects are marked with an asterisk, and indicate the comparison of each level with 0 ECEs (*** = p < 0.001, ** = p < 0.01, * = p < 0.05).

The impact of 4+ days of extreme cold on survival probability was significantly stronger during the nestling stage (β = - 0.56 ± 0.157, z = -3.597, p < 0.001). Survival probability was predicted to decrease by >40%, from 7.7% for individuals that did not experience any cold ECEs to 4.5% for those exposed to four or more cold ECEs (Fig. 4D). Rain ECEs during both the stages, especially during early development, had negligible effects on apparent survival (p>0.1, Fig. 4E,4F). However, a marginally significant negative effect was observed for individuals that experienced three or more days of extreme rain (β = -0.275 ± 0.128, z = -2.145, p < 0.05, Fig. 4F).

Experiencing hot ECEs during both the hatchling and nestling stages, however, appeared to confer a survival advantage, with models predicting significant positive associations between exposure to hot days during development and recruitment probability. For hatchlings, survival probability increased significantly with exposure to hot ECEs, with a 25% increase for individuals experiencing two or more extreme heat events (β = 0.345 ± 0.094, z = 3.649, p < 0.001). The effect was strongest for those exposed to five or more hot ECEs, where survival probability showed a notable 50% increase (β = 0.522 ± 0.15, z = 3.474, p < 0.001, Fig. 4A). Similarly, for nestlings, survival probability increased progressively with higher frequencies of hot ECEs. Individuals exposed to five or more extreme heat events during this stage exhibited the strongest positive association, with recruitment likelihood increasing by 85.7% (β = 0.712 ± 0.121, z = 5.883, p < 0.001, Fig. 4B).

However, when lay date was included in the models, the effects of all ECEs (hot, cold, and rain) on recruitment probability were attenuated, with most estimates decreasing in magnitude and several effects losing statistical significance (Supplementary Table 6, Fig. 4).

## Discussion

With observed climate patterns increasingly aligning with long-term projections, and with extreme climatic events (ECEs) becoming more frequent and intense (IPCC, 2023), understanding their ecological impacts has become a growing priority. Our study provides a comprehensive analysis of the impacts of ECEs during sensitive developmental phases on the growth of great tit nestlings, and their subsequent survival to adulthood, offering critical insights into how climate variability can influence natural populations.

We found that the hatchling stage (0-7 days post hatching) exhibited heightened sensitivity to temperature changes compared to the nestling stage (8-15 days post hatching), with a narrower optimal thermal range, likely due to limited thermoregulatory capability. During early development, hatchlings lack well-developed feathers, leaving them poorly equipped to maintain thermal stability. Exposure to extreme cold during this period was also particularly detrimental to growth. During prolonged extreme cold conditions, hatchlings likely allocate a disproportionate amount of energy towards thermoregulation rather than growth and development (Dawson et al., 2005). Furthermore, increased parental brooding effort, albeit crucial for hatchling survival in such situations, may instead limit food availability due to reduced foraging activity (Rodríguez & Barba, 2016a).

The nestling stage, on the other hand, marked with increased energetic demands of the growing chicks, was more strongly influenced by precipitation patterns (both average and extreme) and extreme heat events. While rain ECEs and elevated average rainfall reduced fledging weight, hot ECEs enhanced growth—a contrast likely mediated by their opposing indirect effects on food availability. Heavy rainfall can deter birds from foraging and can also dislodge caterpillars and other insect prey from vegetation, making them harder to spot by parents (Radford et al., 2001). Conversely, heat can increase insect activity and visibility, potentially boosting prey availability (Mccarty & Winkler, 1999; Schöll et al., 2016). Furthermore, if nestlings are sufficiently warm, parents can dedicate more time to foraging and provisioning, rather than brooding (Dawson et al., 2005; Rodríguez & Barba, 2016b). Although extreme heat can lead to heat stress and dehydration in nestlings, it is important to consider the temperate environment they are developing in. Cooler baseline temperatures would allow nestlings to capitalise on reduced thermoregulatory costs (Andreasson et al., 2018), and benefit from the prey abundance due to hot conditions, promoting their rapid growth (Dawson et al., 2005). Additionally, great tit nestlings primarily consume caterpillars, which have high water content, further reducing their risks of dehydration (Andreasson et al., 2018). These habitat-specific dynamics illustrate how beneficial hot ECEs in temperate climates may contrast sharply with their detrimental impacts in hotter regions (Rodríguez & Barba, 2016b).

Possibly, this temperate climate context may also help explain why our analyses of the rarest and most extreme events, defined by the 1% tails of temperature and rainfall distributions, showed generally weaker and mostly non-significant effects for temperature extremes. Despite such high thresholds, the maximum daily temperature for a 1% hot ECE was around 28°C. While this would certainly qualify as a heatwave, its ecological impact may be modest compared to tropical or arid systems. In contrast, very extreme rainfall events, though also rare, represent more severe deviations from typical conditions in our study area (with peak rainfall reaching up to 40 mm), and were linked to reductions in fledging mass across developmental stages, albeit with smaller effect sizes than the more frequent 5% rain ECEs. However, the extreme rarity of these 1% events inherently limits statistical power, making it difficult to reliably detect their impacts even with a robust dataset.

Apart from isolated effects we found that ECEs can interact with ambient environmental conditions to collectively influence chick growth outcomes. During the hatchling stage, while rain and hot ECEs did not directly affect growth, their interactions with mean climatic variables proved significant. The combination of extreme rainfall and higher average temperatures, or the converse of extreme heat combined with consistently higher precipitation, predicted a particularly challenging scenario for early development. These synergistic effects likely stem from the limited thermoregulatory capabilities of young hatchlings, exacerbated by the resource limitations imposed by prey unavailability and altered parental foraging behaviour during extreme rainfall. Interestingly, nestlings showed a contrasting pattern, where the negative effect of higher rainfall was reversed in the presence of extreme heat, although this interaction was marginally significant (Supplementary Table 4). This unexpected result suggests that older nestlings might be better equipped to handle, and even benefit from, the combination of warmth and moisture, unlike hatchlings, which were predicted to be especially vulnerable to compounded climatic stressors.

Perhaps most importantly, our findings further suggested that the consequences of ECEs on chick growth outcomes are highly contingent upon seasonal breeding phenology. Being part of early clutches confers significant advantages, as abundant food resources earlier in the season may support better growth outcomes. Furthermore, earlier laying allows parents greater control over incubation timing, enabling them to match hatching more closely with the caterpillar peak and optimise provisioning opportunities (Simmonds et al., 2017). Consequently, our models indicated that broods laid earlier in the season demonstrate a degree of resilience or even benefit from hot ECEs during the hatchling stage, whereas late broods face significantly greater challenges. Hot ECEs during early development exacerbated the disadvantages of being part of a late brood, where reduced food availability combined with high temperatures may be particularly detrimental to growth. Similarly, rain ECEs during the nestling stage intensified resource limitations for late-clutch offspring, by further reducing foraging opportunities for already scarce prey late in the season.

While ECEs can exacerbate poor resource availability and challenging climatic conditions, their impacts in isolation are not extremely strong in this population. As our results show, nestlings from early broods may even benefit from higher temperatures, experiencing them as moderate warmth rather than extreme heat. However, while breeding earlier within a season may help offset the negative impacts of ECEs, longer-term shifts in breeding phenology add further complexity to population responses. Over recent decades, our study population has shifted to earlier breeding, likely as an adaptive response to warming temperatures (Charmantier et al., 2008). This has inadvertently increased exposure to extreme cold events early in the breeding season (Regan & Sheldon, 2023), which our findings show to be damaging to nestling growth. Consequently, although earlier breeding within a season may mitigate some challenges, the overarching trend toward earlier laying could increase the population’s susceptibility to detrimental cold extremes. This suggests that adaptive shifts in phenology may not entirely eliminate the risks posed by evolving extreme climatic conditions, emphasising the need for continuous monitoring to assess how future climate scenarios affect population resilience.

Our analysis also revealed weak long-term effects of ECEs, with extreme cold and rain during the nestling period reducing the likelihood of recruiting to the breeding population, likely due to thermal stress and resource depletion. Previous studies have also documented the negative impacts of increased rainfall during critical developmental periods on offspring survival (Arct et al., 2025; Pipoly et al., 2013; Schöll & Hille, 2020). Conversely, survival to adulthood was predicted to be more likely if chicks experienced hot ECEs throughout their development. These findings align with an experimental study on blue tits with artificially elevated nest temperatures (Andreasson et al., 2018). Higher ambient temperatures during the nestling stage were shown to be associated with a higher number of recruits in a wild population of collared flycatchers (Ficedula albicollis) in a recent study (Arct et al., 2025). However, no effects of heat treatments on post-fledging survival were observed in another experiment with great tits (Rodríguez & Barba, 2016b). Accounting for the relative egg-laying timing in our models diminished the effects of ECEs on apparent survival, suggesting that selection pressures favouring earlier clutches may have enhanced resilience to long-term ECE impacts. However, while earlier breeding may have buffered this population against some climatic challenges, it may not always be protective as global temperatures continue to rise. For example, while hot ECEs currently have limited negative impacts in this temperate population–likely because they do not exceed thermal thresholds (Rodríguez & Barba, 2016b)–future increases in heat intensity could impose significant stress on chicks.

Despite these insights, our study has limitations that warrant consideration. First, our definition of ECEs assumes that their effects have remained constant over a 60-year period. Shifting climate norms and increased variability mean that past ‘extreme’ events may now be more common, potentially altering their ecological significance over time. Second, we focused on fledging mass and recruitment probability as key metrics but lacked data on intermediate developmental stages where compensatory mechanisms might occur. Furthermore, our study utilised broad-scale weather data, assuming uniform ambient climatic conditions across all nests, which overlooks the possible effect of fine-scale habitat heterogeneity. Fine-scale microenvironmental variation could lead to different ECE exposures and impacts, even for broods in relatively close proximity. On the same note, while our study identified significant impacts of precipitation on chick growth, we relied on daily rainfall data that may not capture fine-scale temporal dynamics. Consecutive days of extreme rain likely have different ecological consequences than intermittent heavy rainfall events. Future studies should incorporate high-resolution rainfall data to better understand how temporal patterns influence food availability and parental behaviour.

Finally, it is important to consider the broader implications of our findings for other populations. Are the patterns we observed in Wytham Woods generalizable, or are they specific to this population due to its unique environmental context and evolutionary history? Understanding the consistency of these responses across different populations and species is crucial for predicting the wider impacts of climate change and identifying the factors that promote or constrain adaptation. Our findings contribute to this understanding by emphasizing the complexity of ecological responses, the importance of developmental stage-specific vulnerabilities, and the role of phenological shifts in mitigating climate impacts. Future research should therefore focus on incorporating finer-scale environmental data, investigating the physiological mechanisms underlying the observed responses, and exploring how microclimates and individual variation buffer against extreme events. Ultimately, as ECEs become more frequent and severe, understanding these dynamics will be essential for informing conservation strategies and predicting population-level responses in a rapidly changing world.

## Acknowledgements

We are grateful to the numerous people who helped collect long-term data for the Wytham tit study over the past 75 years. The long-term population study has been supported by numerous funding sources, including recently by grants from BBSRC (BB/L006081/1) and NERC (NE/K006274/1, NE/S010335/1). Devi Satarkar is supported by the Oxford-Oxitec Graduate Scholarship. Irem Sepil is supported by a Royal Society Dorothy Hodgkin Fellowship (DHF\R1\211084).

## Author contributions

All authors contributed to the ideas of this study. Devi Satarkar conducted the statistical analyses with substantial inputs on methodology from David López-Idiáquez and Ben C. Sheldon. Devi Satarkar drafted the manuscript with David López-Idiáquez, Irem Sepil and Ben C. Sheldon providing critical feedback. All authors approved the final manuscript.

## Data and code availability

Data and code to reproduce all analyses are available at https://github.com/devisatarkar/ECEchickweight_great-tits

