## Supplementary Information for "Developmental stage-specific responses to extreme climatic events and environmental variability in great tit nestlings"

**Table 1. Ambient climate models:** Outputs of linear mixed models for fledging mass with ambient climate measures (mean temperature/ mean rainfall) during specific developmental stages (hatchling/nestling) as explanatory variables along with laydate, and clutch size as fixed effects. Year of birth, brood identity, mother identity and natal nest box are included as random effects. All fixed effects are scaled to a mean of zero and standard deviation of one. Significant terms (*p <0.05)* are in bold.

| Variable | Estimate | Std. Error | t | p |
|---|---|---|---|---|
| *Average temperature (hatchling stage)* | | | | |
| Mean temperature | 0.1202 | 0.01994 | 6.03 | **< 0.001** |
| **(Mean temperature)$^2$** | **-0.0763** | 0.01099 | -6.94 | **< 0.001** |
| Lay date | -0.3343 | 0.02021 | -16.54 | **< 0.001** |
| Clutch size | -0.2474 | 0.01409 | -17.55 | **< 0.001** |
| *Average temperature (hatchling stage) – using splines* | | | | |
| **Mean temperature.1** | **0.8245** | 0.1548 | 5.33 | **< 0.001** |
| **Mean temperature.2** | **0.6015** | 0.1795 | 3.35 | **0.000808** |
| **Mean temperature.3** | **0.7902** | 0.1262 | 6.26 | **< 0.001** |
| **Mean temperature.4** | **0.7237** | 0.3826 | 1.89 | **0.0586** |
| Mean temperature.5 | 0.03598 | 0.2200 | 0.16 | 0.8701 |
| Lay date | -0.3317 | 0.02026 | -16.37 | **< 0.001** |
| Clutch size | -0.2477 | 0.01410 | -17.57 | **< 0.001** |
| *Average temperature (nestling stage)* | | | | |
| Mean temperature | 0.1344 | 0.01812 | 7.42 | **< 0.001** |
| **(Mean temperature)$^2$** | **-0.04692** | 0.01009 | -4.65 | **< 0.001** |
| Lay date | -0.3515 | 0.02039 | -17.24 | **< 0.001** |
| Clutch size | -0.2485 | 0.01412 | -17.61 | **< 0.001** |
| *Average temperature (nestling stage) – using splines* | | | | |

| | | | | |
|---|---|---|---|---|
| **Mean temperature.1** | **0.5032** | 0.1848 | 2.72 | **0.00649** |
| **Mean temperature.2** | **0.3726** | 0.2049 | 1.82 | **0.06905** |
| **Mean temperature.3** | **0.8374** | 0.1546 | 5.42 | **< 0.001** |
| Mean temperature.4 | 0.09307 | 0.4643 | 0.20 | 0.84114 |
| Mean temperature.5 | -0.3220 | 0.4166 | -0.77 | 0.43955 |
| Lay date | -0.3516 | 0.02047 | -17.17 | **< 0.001** |
| Clutch size | -0.2488 | 0.01412 | -17.63 | **< 0.001** |
| *Average rainfall (hatchling stage)* | | | | |
| **Mean rainfall** | **-0.06288** | 0.01813 | -3.47 | **0.000526** |
| Lay date | -0.2918 | 0.01871 | -15.59 | **< 0.001** |
| Clutch size | -0.2387 | 0.01408 | -16.96 | **< 0.001** |
| *Average rainfall (nestling stage)* | | | | |
| **Mean rainfall** | **-0.1439** | 0.01580 | -9.11 | **< 0.001** |
| Lay date | -0.2857 | 0.01867 | -15.30 | **< 0.001** |
| Clutch size | -0.2349 | 0.01404 | -16.72 | **< 0.001** |

**Table 2. Extreme climate events (frequency) models:** Outputs of linear mixed models for fledging mass with number of ECEs during specific developmental stages (hatchling/nestling) as explanatory variables along with mean temperature during the relevant stage, laydate, and clutch size as fixed effects. Year of birth, brood identity, mother identity and natal nest box are included as random effects. All fixed effects are scaled to a mean of zero and standard deviation of one. Significant terms (*p <0.05)* are in bold. Here, ECEs are calculated with a 5% threshold.

| Variable | Estimate | Std. Error | t | p |
|---|---|---|---|---|
| *Number of hot ECEs (hatchling stage)* | | | | |
| Mean temperature | 0.1051 | 0.02749 | 3.83 | **< 0.001** |
| Lay date | -0.3332 | 0.02144 | -15.54 | **< 0.001** |

| | | | | |
|---|---|---|---|---|
| Clutch size | -0.2445 | 0.01416 | -17.27 | **< 0.001** |
| Number of hot ECEs | -0.02235 | 0.02258 | -0.99 | 0.3222 |
| *Number of hot ECEs (nestling stage)* | | | | |
| Mean temperature | 0.0614 | 0.0237 | 2.59 | **0.009** |
| Lay date | -0.3202 | 0.0216 | -14.82 | **< 0.001** |
| Clutch size | -0.2444 | 0.0142 | -17.25 | **< 0.001** |
| **Number of hot ECEs** | **0.0914** | 0.0240 | 3.81 | **0.00014** |
| *Number of hot ECEs (nestling stage) – using splines* | | | | |
| Number of hot ECEs.1 | 0.08188 | 0.07484 | 1.09 | 0.274 |
| **Number of hot ECEs.2** | **-0.8576** | 0.2392 | -3.59 | **< 0.001** |
| **Number of hot ECEs.3** | **1.883** | 0.4019 | 4.68 | **< 0.001** |
| Mean temperature | 0.07136 | 0.02424 | 2.94 | **0.003** |
| Lay date | -0.3236 | 0.02176 | -14.87 | **< 0.001** |
| Clutch size | -0.2447 | 0.01417 | -17.27 | **< 0.001** |
| *Number of cold ECEs (hatchling stage)* | | | | |
| Mean temperature | 0.05067 | 0.02205 | 2.30 | **0.0216** |
| Lay date | -0.3224 | 0.02024 | -15.93 | **< 0.001** |
| Clutch size | -0.2428 | 0.01411 | -17.22 | **< 0.001** |
| **Number of cold ECEs** | **-0.06948** | 0.02089 | -3.33 | **0.00089** |
| *Number of cold ECEs (hatchling stage) – using splines* | | | | |
| **Number of cold ECEs.1** | **0.3557** | 0.1077 | 3.30 | **0.00096** |
| **Number of cold ECEs.2** | **-1.008** | 0.3647 | -2.77 | **0.00571** |
| Number of cold ECEs.3 | 0.2781 | 0.5891 | 0.47 | 0.637 |
| Mean temperature | 0.05492 | 0.02216 | 2.48 | **0.0132** |
| Lay date | -0.3304 | 0.02033 | -16.25 | **< 0.001** |

| | | | | |
|---|---|---|---|---|
| Clutch size | -0.2436 | 0.01410 | -17.28 | **< 0.001** |
| *Number of cold ECEs (nestling stage)* | | | | |
| Mean temperature | 0.1734 | 0.02080 | 8.34 | **< 0.001** |
| Lay date | -0.3705 | 0.02088 | -17.74 | **< 0.001** |
| Clutch size | -0.2524 | 0.01414 | -17.85 | **< 0.001** |
| Number of cold ECEs | 0.09059 | 0.01833 | 4.97 | **< 0.001** |
| *Number of rain ECEs (hatchling stage)* | | | | |
| Mean temperature | 0.08192 | 0.01954 | 4.19 | **< 0.001** |
| Lay date | -0.3243 | 0.02026 | -16.01 | **< 0.001** |
| Clutch size | -0.2431 | 0.01411 | -17.23 | **< 0.001** |
| Number of rain ECEs | -0.02431 | 0.01637 | -1.49 | 0.137 |
| *Number of rain ECEs (nestling stage)* | | | | |
| Mean temperature | 0.1095 | 0.01805 | 6.06 | **< 0.001** |
| Lay date | -0.3367 | 0.02051 | -16.42 | **< 0.001** |
| Clutch size | -0.2451 | 0.01414 | -17.33 | **< 0.001** |
| **Number of rain ECEs** | **-0.07346** | 0.01619 | -4.54 | **< 0.001** |

**Table 3. Extreme climate events (binary) models:** Outputs of linear mixed models for fledging mass with presence of at least 1 ECE (1% or 5%) during specific developmental stages (hatchling/nestling) as explanatory variables along with mean temperature during the relevant stage, laydate, and clutch size as fixed effects. Year of birth, brood identity, mother identity and natal nest box are included as random effects. All fixed effects are scaled to a mean of zero and standard deviation of one. Significant terms (*p <0.05)* are in bold.

| Variable | β | Std. Error | t | p | β | Std. Error | t | p |
|---|---|---|---|---|---|---|---|---|
| *Presence of at least 1 hot (1%) ECE (**hatchling**)* | | | | | *Presence of at least 1 hot (5%) ECE (**hatchling**)* | | | |

| | | | | | | | | |
|---|---|---|---|---|---|---|---|---|
| Presence of hot ECE | -0.014 | 0.018 | -0.797 | 0.425 | 0.047 | 0.020 | 2.319 | **0.0204** |
| Mean temperature | 0.095 | 0.023 | 4.221 | **< 0.001** | 0.048 | 0.025 | 1.866 | 0.0621 |
| Clutch size | -0.244 | 0.014 | -17.260 | **< 0.001** | -0.241 | 0.014 | -17.089 | **< 0.001** |
| Lay date | -0.329 | 0.020 | -16.054 | **< 0.001** | -0.314 | 0.021 | -14.980 | **< 0.001** |
| *Presence of at least 1 hot (1%) ECE (**nestling**)* | | | | | *Presence of at least 1 hot (5%) ECE (**nestling**)* | | | |
| Presence of hot ECE | 0.023 | 0.020 | 1.138 | 0.255 | 0.012 | 0.020 | 0.597 | 0.551 |
| Mean temperature | 0.110 | 0.020 | 5.479 | **< 0.001** | 0.113 | 0.022 | 5.18 | **< 0.001** |
| Clutch size | -0.248 | 0.014 | -17.533 | **< 0.001** | -0.248 | 0.014 | -17.547 | **< 0.001** |
| Lay date | -0.343 | 0.021 | -16.539 | **< 0.001** | -0.344 | 0.021 | -16.164 | **< 0.001** |
| *Presence of at least 1 cold (1%) ECE (**hatchling**)* | | | | | *Presence of at least 1 cold (5%) ECE (**hatchling**)* | | | |
| Presence of cold ECE | 0.013 | 0.013 | 1.004 | 0.315 | -0.031 | 0.020 | -1.582 | 0.113785 |
| Mean temperature | 0.088 | 0.019 | 4.504 | **< 0.001** | 0.071 | 0.021 | 3.339 | **< 0.001** |
| Clutch size | -0.244 | 0.014 | -17.259 | **< 0.001** | -0.243 | 0.014 | -17.239 | **< 0.001** |
| Lay date | -0.328 | 0.020 | -16.158 | **< 0.001** | -0.324 | 0.020 | -15.982 | **< 0.001** |
| *Presence of at least 1 cold (1%) ECE (**nestling**)* | | | | | *Presence of at least 1 cold (5%) ECE (**nestling**)* | | | |
| Presence of cold ECE | **0.087** | 0.017 | 4.983 | **< 0.001** | **0.096** | 0.017 | 5.667 | **< 0.001** |
| Mean temperature | 0.132 | 0.018 | 7.333 | **< 0.001** | 0.174 | 0.020 | 8.614 | **< 0.001** |
| Clutch size | -0.249 | 0.014 | -17.662 | **< 0.001** | -0.251 | 0.014 | -17.776 | **< 0.001** |
| Lay date | -0.356 | 0.020 | -17.416 | **< 0.001** | -0.369 | 0.021 | -17.817 | **< 0.001** |
| *Presence of at least 1 rain (1%) ECE (**hatchling**)* | | | | | *Presence of at least 1 rain (5%) ECE (**hatchling**)* | | | |

| | | | | | | | | |
|---|---|---|---|---|---|---|---|---|
| Presence of rain ECE | **-0.041** | 0.016 | -2.578 | **0.00995** | -0.012 | 0.016 | -0.751 | 0.453 |
| Mean temperature | 0.084 | 0.019 | 4.324 | **< 0.001** | 0.084 | 0.020 | 4.293 | **< 0.001** |
| Clutch size | -0.243 | 0.014 | -17.235 | **< 0.001** | -0.243 | 0.014 | -17.242 | **< 0.001** |
| Lay date | -0.325 | 0.020 | -16.065 | **< 0.001** | -0.325 | 0.020 | -16.058 | **< 0.001** |
| *Presence of at least 1 rain (1%) ECE (**nestling**)* | | | | | *Presence of at least 1 rain (5%) ECE (**nestling**)* | | | |
| Presence of rain ECE | **-0.042** | 0.017 | -2.488 | **0.0129** | **-0.077** | 0.016 | -4.86 | **< 0.001** |
| Mean temperature | 0.115 | 0.018 | 6.364 | **< 0.001** | 0.105 | 0.018 | 5.786 | **< 0.001** |
| Clutch size | -0.248 | 0.014 | -17.554 | **< 0.001** | -0.246 | 0.014 | -17.39 | **< 0.001** |
| Lay date | -0.343 | 0.020 | -16.773 | **< 0.001** | -0.341 | 0.020 | -16.7 | **< 0.001** |

**Table 4. Interaction models (ambient climate x ECE frequency):** Outputs of linear mixed models for fledging mass with number of ECEs during specific developmental stages (hatchling/nestling) interacting with relevant ambient climate measures as predictors, along with laydate and clutch size as fixed effects. Year of birth, brood identity, mother identity and natal nest box are included as random effects. All fixed effects are scaled to a mean of zero and standard deviation of one. Significant terms (*p <0.05)* are in bold. Here, ECEs are calculated with a 5% threshold.

| Variable | Estimate | Std. Error | t | p |
|---|---|---|---|---|
| *Average temperature x number of rain ECEs (hatchling stage)* | | | | |
| Mean temperature | 0.10280 | 0.02055 | 5.000 | **< 0.001** |
| (Mean temperature)$^2$ | -0.08914 | 0.01140 | -7.816 | **< 0.001** |
| Number of rain ECEs | -0.008569 | 0.01932 | -0.443 | 0.657 |
| **Number of rain ECEs x mean temp** | **-0.08596** | 0.01729 | -4.973 | **< 0.001** |

| | | | | |
|---|---|---|---|---|
| Number of rain ECEs x (mean temp)$^2$ | -0.01853 | 0.01240 | -1.494 | 0.135 |
| Lay date | -0.32330 | 0.02031 | -15.915 | **< 0.001** |
| Clutch size | -0.24490 | 0.01408 | -17.391 | **< 0.001** |
| *Average temperature x number of rain ECEs (nestling stage)* | | | | |
| Mean temperature | 0.12780 | 0.01871 | 6.831 | **< 0.001** |
| (Mean temperature)$^2$ | -0.04408 | 0.01047 | -4.210 | **< 0.001** |
| Number of rain ECEs | **-0.09363** | **0.01879** | -4.983 | **< 0.001** |
| Number of rain ECEs x mean temp | -0.01330 | 0.01887 | -0.705 | 0.4808 |
| Number of rain ECEs x (mean temp)$^2$ | 0.02973 | 0.01329 | 2.236 | 0.0253 |
| Lay date | -0.33750 | 0.02056 | -16.418 | **< 0.001** |
| Clutch size | -0.24470 | 0.01413 | -17.317 | **< 0.001** |
| *Average rainfall x number of hot ECEs (hatchling stage)* | | | | |
| Mean rainfall | -0.08781 | 0.01867 | -4.703 | **< 0.001** |
| Number of hot ECEs | **-0.06939** | 0.02115 | -3.282 | **0.00104** |
| **Number of hot ECEs x mean rainfall** | **-0.16190** | 0.02197 | -7.367 | **< 0.001** |
| Lay date | -0.28250 | 0.01872 | -15.088 | **< 0.001** |
| Clutch size | 0.23510 | 0.01405 | -16.729 | **< 0.001** |
| *Average rainfall x number of hot ECEs (nestling stage)* | | | | |
| Mean rainfall | -0.11430 | 0.01682 | -6.795 | **< 0.001** |
| Number of hot ECEs | **0.12860** | 0.02045 | 6.288 | **< 0.001** |

| | | | | |
|---|---|---|---|---|
| Number of hot ECEs x mean rainfall | 0.05163 | 0.02041 | 2.530 | 0.0114 |
| Lay date | -0.28690 | 0.01864 | -15.389 | **< 0.001** |
| Clutch size | -0.23560 | 0.01403 | -16.800 | **< 0.001** |
| *Average rainfall x number of cold ECEs (hatchling stage)* | | | | |
| Mean rainfall | -0.05072 | 0.01831 | -2.770 | **0.00561** |
| Number of cold ECEs | **-0.08035** | 0.01874 | -4.287 | **< 0.001** |
| Number of cold ECEs x mean rainfall | 0.03919 | 0.01985 | 1.974 | 0.04838 |
| Lay date | -0.30360 | 0.01890 | -16.070 | **< 0.001** |
| Clutch size | -0.24000 | 0.01407 | -17.059 | **< 0.001** |
| *Average rainfall x number of cold ECEs (nestling stage)* | | | | |
| Mean rainfall | -0.14640 | 0.01593 | -9.190 | **< 0.001** |
| Number of cold ECEs | 0.03267 | 0.01587 | 2.059 | 0.03952 |
| Number of cold ECEs x mean rainfall | 0.05829 | 0.01624 | 3.589 | **0.000334** |
| Lay date | -0.28960 | 0.01870 | -15.488 | **< 0.001** |
| Clutch size | -0.23540 | 0.01404 | -16.767 | **< 0.001** |

**Table 5. Interaction models (relative laydate x ECE frequency):** Outputs of linear mixed models for fledging mass with number of ECEs during specific developmental stages (hatchling/nestling) interacting with relative lay date as predictors, along with mean temperature and clutch size as fixed effects. Year of birth, brood identity, mother identity and natal nest box are included as random effects. All fixed effects are scaled to a mean of zero and standard deviation of one. Significant terms (*p <0.05)* are in bold. Here, ECEs are calculated with a 5% threshold.

| Variable | Estimate | Std. Error | t | p |
|---|---|---|---|---|
| *Base model for relative laydate* | | | | |
| Relative laydate | -0.13960 | 0.01443 | -9.676 | **< 0.001** |
| **(Relative laydate)$^2$** | **-0.04080** | 0.00464 | -8.791 | **< 0.001** |
| Clutch size | -0.23560 | 0.01404 | -16.787 | **< 0.001** |
| *Relative laydate x number of hot ECEs (hatchling stage)* | | | | |
| Relative laydate | -0.16190 | 0.01596 | -10.144 | **< 0.001** |
| (Relative laydate)$^2$ | -0.04306 | 0.00465 | -9.268 | **< 0.001** |
| Mean temperature | 0.11280 | 0.02724 | 4.142 | **< 0.001** |
| Number of hot ECEs | -0.02641 | 0.02388 | -1.106 | 0.269 |
| **Number of hot ECEs x relative laydate** | **-0.06710** | 0.01295 | -5.181 | **< 0.001** |
| Number of hot ECEs x (relative laydate)$^2$ | 0.006421 | 0.00493 | 1.304 | 0.192 |
| Clutch size | -0.23970 | 0.01409 | -17.008 | **< 0.001** |
| *Relative laydate x number of hot ECEs (nestling stage)* | | | | |
| Relative laydate | -0.15770 | 0.01646 | -9.586 | **< 0.001** |
| (Relative laydate)$^2$ | -0.03956 | 0.00467 | -8.480 | **< 0.001** |
| Mean temperature | 0.05694 | 0.02365 | 2.407 | 0.01609 |
| Number of hot ECEs | 0.07277 | 0.02432 | 2.992 | **0.00278** |
| Number of hot ECEs x relative laydate | 0.01174 | 0.01738 | 0.676 | 0.49926 |
| Number of hot ECEs x (relative laydate)$^2$ | 0.00822 | 0.00490 | 1.677 | 0.09364 |

| | | | | |
|---|---|---|---|---|
| Clutch size | -0.24110 | 0.01414 | -17.049 | **< 0.001** |
| *Relative laydate x number of cold ECEs (hatchling stage)* | | | | |
| Relative laydate | -0.1575 | 0.01522 | -10.349 | **< 0.001** |
| (Relative laydate)$^2$ | -0.04095 | 0.00501 | -8.173 | **< 0.001** |
| Mean temperature | 0.07051 | 0.02191 | 3.218 | **0.0013** |
| Number of cold ECEs | -0.01598 | 0.02347 | -0.681 | 0.4960 |
| Number of cold ECEs x relative laydate | **0.1000** | 0.01585 | 6.313 | **< 0.001** |
| Number of cold ECEs x (relative laydate)$^2$ | -0.01439 | 0.00772 | -1.864 | 0.0623 |
| Clutch size | -0.2368 | 0.01404 | -16.861 | **< 0.001** |
| *Relative laydate x number of cold ECEs (nestling stage)* | | | | |
| Relative laydate | -0.1964 | 0.01594 | -12.321 | **< 0.001** |
| (Relative laydate)$^2$ | -0.0441 | 0.00497 | -8.881 | **< 0.001** |
| Mean temperature | 0.1621 | 0.02069 | 7.834 | **< 0.001** |
| Number of cold ECEs | 0.1432 | 0.02270 | 6.305 | **< 0.001** |
| Number of cold ECEs x relative laydate | **0.0377** | 0.01279 | 2.945 | **0.00324** |
| Number of cold ECEs x (relative laydate)$^2$ | **-0.0394** | 0.00885 | -4.449 | **< 0.001** |
| Clutch size | -0.2491 | 0.01409 | -17.682 | **< 0.001** |
| *Relative laydate x number of rain ECEs (hatchling stage)* | | | | |
| Relative laydate | -0.1643 | 0.01522 | -10.796 | **< 0.001** |
| (Relative laydate)$^2$ | -0.04215 | 0.00464 | -9.076 | **< 0.001** |

| | | | | |
|---|---|---|---|---|
| Mean temperature | 0.09277 | 0.01939 | 4.784 | **< 0.001** |
| Number of rain ECEs | 0.00335 | 0.01818 | 0.184 | 0.854 |
| Number of rain ECEs x relative laydate | 0.06374 | 0.01318 | 4.835 | **< 0.001** |
| Number of rain ECEs x (relative laydate)$^2$ | -0.01935 | 0.00481 | -4.020 | **< 0.001** |
| Clutch size | -0.2414 | 0.01404 | -17.193 | **< 0.001** |
| *Relative laydate x number of rain ECEs (nestling stage)* | | | | |
| Relative laydate | -0.1650 | 0.01582 | -10.433 | **< 0.001** |
| (Relative laydate)$^2$ | -0.04319 | 0.00479 | -9.024 | **< 0.001** |
| Mean temperature | 0.09695 | 0.01805 | 5.372 | **< 0.001** |
| Number of rain ECEs | **-0.05820** | 0.01751 | -3.323 | **0.000893** |
| Number of rain ECEs x relative laydate | -0.01501 | 0.01628 | -0.922 | 0.356464 |
| **Number of rain ECEs x (relative laydate)$^2$** | **-0.01414** | 0.00606 | -2.334 | **0.019614** |
| Clutch size | -0.2411 | 0.01410 | -17.100 | **< 0.001** |

**Table 6. Local recruitment models:** Outputs of generalised linear mixed models for recruitment probability with number of ECEs during specific developmental stages (hatchling/nestling) as predictors, along with clutch size as fixed effect. Outputs of additional models with laydate as fixed effect have also been detailed below. ECEs are treated here as categorical variables. Year of birth, brood identity, mother identity and natal nest box are included as random effects. All fixed effects are scaled to a mean of zero and standard deviation of one. Significant terms (*p <0.05)* are in bold. Here, ECEs are calculated with a 5% threshold.

| Variable | β | Std. Error | z | p | β | Std. Error | z | p |
|---|---|---|---|---|---|---|---|---|

| | | | | | | | | |
|---|---|---|---|---|---|---|---|---|
| *Number of Cold ECEs (hatchling stage)* | | | | | *with laydate* | | | |
| **1 Cold ECE** | **-0.2418** | 0.07388 | -3.273 | **0.00106** | -0.14016 | 0.07507 | -1.867 | 0.0619 |
| 2 Cold ECEs | -0.02525 | 0.08889 | -0.284 | 0.77642 | -0.03618 | 0.08958 | -0.404 | 0.6863 |
| **3 Cold ECEs** | **-0.26187** | 0.10499 | -2.494 | **0.01262** | **-0.22566** | 0.10568 | -2.135 | **0.0327** |
| **4+ Cold ECEs** | **-0.30369** | 0.12826 | -2.368 | **0.01789** | -0.20996 | 0.13088 | -1.604 | 0.1087 |
| Mean temperature | -0.14797 | 0.02337 | -6.332 | **< 0.001** | -0.03616 | 0.02486 | -1.454 | 0.1459 |
| Clutch size | -0.04323 | 0.01513 | -2.858 | **0.00427** | -0.1079 | 0.01591 | -6.783 | **< 0.001** |
| Lay date | – | – | – | **–** | -0.33045 | 0.02486 | -13.295 | **< 0.001** |
| *Number of Cold ECEs (nestling stage)* | | | | | *with laydate* | | | |
| **1 Cold ECE** | **-0.15801** | 0.06662 | -2.372 | **0.017695** | -0.02927 | 0.0683 | -0.429 | 0.6683 |
| 2 Cold ECEs | 0.041865 | 0.13519 | 0.31 | 0.756821 | 0.1619 | 0.13507 | 1.199 | 0.2307 |
| 3 Cold ECEs | 0.004844 | 0.12191 | 0.04 | 0.968308 | 0.19524 | 0.12305 | 1.587 | 0.1126 |
| **4+ Cold ECEs** | **-0.56687** | 0.15759 | -3.597 | **0.000322** | -0.16141 | 0.16179 | -0.998 | 0.3184 |
| Mean temperature | -0.17697 | 0.02122 | -8.337 | **< 0.001** | -0.04842 | 0.02408 | -2.011 | **0.0444** |
| Clutch size | -0.04353 | 0.01513 | -2.878 | **0.004008** | -0.10503 | 0.01599 | -6.569 | **< 0.001** |
| Lay date | – | – | – | **–** | -0.31035 | 0.0259 | -11.984 | **< 0.001** |
| *Number of Rain ECEs (hatchling stage)* | | | | | *with laydate* | | | |
| 1 Rain ECE | -0.04466 | 0.04182 | -1.068 | 0.2856 | -0.005418 | 0.042032 | -0.129 | 0.897 |
| 2 Rain ECEs | -0.07559 | 0.06781 | -1.115 | 0.2649 | -0.035606 | 0.068786 | -0.518 | 0.605 |
| 3+ Rain ECEs | 0.13694 | 0.19798 | 0.692 | 0.4891 | 0.120993 | 0.205366 | 0.589 | 0.556 |
| Mean temperature | -0.12121 | 0.02063 | -5.874 | **< 0.001** | -0.011484 | 0.022206 | -0.517 | 0.605 |
| Clutch size | -0.04333 | 0.01513 | -2.863 | **0.0042** | -0.110118 | 0.015911 | -6.921 | **< 0.001** |

| | | | | | | | | |
|---|---|---|---|---|---|---|---|---|
| Lay date | – | – | – | **–** | -0.33384 | 0.024779 | -13.473 | **< 0.001** |
| *Number of Rain ECEs (nestling stage)* | | | | | *with laydate* | | | |
| 1 Rain ECE | -0.002987 | 0.042626 | -0.07 | 0.9441 | -0.016176 | 0.042829 | -0.378 | 0.7057 |
| 2 Rain ECEs | -0.079729 | 0.062284 | -1.28 | 0.2005 | -0.001325 | 0.063122 | -0.021 | 0.9833 |
| **3+ Rain ECEs** | **-0.275035** | 0.128208 | -2.145 | **0.0319** | -0.114075 | 0.129947 | -0.878 | 0.38 |
| Mean temperature | -0.149838 | 0.01889 | -7.932 | **< 0.001** | -0.051995 | 0.020788 | -2.501 | **0.0124** |
| Clutch size | -0.040311 | 0.015127 | -2.665 | **0.0077** | -0.105978 | 0.015981 | -6.632 | **< 0.001** |
| Lay date | – | – | – | **–** | -0.313345 | 0.025328 | -12.371 | **< 0.001** |
| *Number of Hot ECEs (hatchling stage)* | | | | | *with laydate* | | | |
| **1 Hot ECE** | **0.12491** | 0.0626 | 1.995 | **0.046018** | -0.022391 | 0.064208 | -0.349 | 0.727 |
| **2 Hot ECEs** | **0.34596** | 0.0948 | 3.649 | **0.000263** | 0.031739 | 0.098806 | 0.321 | 0.748 |
| **3 Hot ECEs** | **0.24191** | 0.10574 | 2.288 | **0.022148** | -0.164768 | 0.111179 | -1.482 | 0.138 |
| **4 Hot ECEs** | **0.46152** | 0.1761 | 2.621 | **0.008771** | 0.088114 | 0.180568 | 0.488 | 0.626 |
| **5+ Hot ECEs** | **0.52201** | 0.15026 | 3.474 | **0.000513** | 0.029637 | 0.156089 | 0.19 | 0.849 |
| Mean temperature | -0.19485 | 0.02826 | -6.895 | **< 0.001** | -0.001021 | 0.032092 | -0.032 | 0.975 |
| Clutch size | -0.04543 | 0.01514 | -3.002 | **0.002686** | -0.111208 | 0.015944 | -6.975 | **< 0.001** |
| Lay date | – | – | – | **–** | -0.338837 | 0.025942 | -13.061 | **< 0.001** |
| *Number of Hot ECEs (nestling stage)* | | | | | *with laydate* | | | |
| **1 Hot ECE** | **0.23385** | 0.05376 | 4.35 | **< 0.001** | 0.08144 | 0.05566 | 1.463 | 0.143399 |
| **2 Hot ECEs** | **0.34338** | 0.08938 | 3.842 | **0.000122** | 0.1952 | 0.09084 | 2.149 | **0.031642** |
| **3 Hot ECEs** | **0.52889** | 0.10655 | 4.964 | **< 0.001** | 0.23387 | 0.11032 | 2.12 | **0.034011** |

| | | | | | | | | |
|---|---|---|---|---|---|---|---|---|
| **4 Hot ECEs** | **0.55928** | 0.11811 | 4.735 | **< 0.001** | 0.23986 | 0.12172 | 1.971 | **0.048772** |
| **5+ Hot ECEs** | **0.7122** | 0.12106 | 5.883 | **< 0.001** | 0.27359 | 0.12834 | 2.132 | **0.033026** |
| Mean temperature | -0.25472 | 0.02409 | -10.572 | **< 0.001** | -0.09981 | 0.02776 | -3.595 | **0.000324** |
| Clutch size | -0.04643 | 0.01515 | -3.065 | **0.002174** | -0.10275 | 0.01602 | -6.416 | **< 0.001** |
| Lay date | – | – | – | **–** | -0.29293 | 0.02654 | -11.037 | **< 0.001** |

**Table 7.** Higher frequencies of ECEs were grouped into a single categorical level, due to the reduced number of individuals experiencing high frequencies of ECEs. Local recruitment models from Table 6 used categorical ECE variables for analysis. Blue shaded areas indicate the values that were combined for each type of ECE in each developmental period.

| **ECE type** | **Frequency** | | | | | | | |
|---|---|---|---|---|---|---|---|---|
| | 0 | 1 | 2 | 3 | 4 | 5 | 6 | 7 |
| **Hatchling** | | | | | | | | |
| Hot | 68167 | 8084 | 2568 | 3262 | 595 | 633 | 565 | 61 |
| Cold | 70622 | 5036 | 4015 | 2382 | 1443 | 424 | 13 | 0 |
| Rain | 59522 | 17499 | 6432 | 459 | 23 | 0 | 0 | 0 |
| **Nestling** | | | | | | | | |
| Hot | 60344 | 12561 | 3821 | 2537 | 2055 | 1594 | 934 | 84 |
| Cold | 74884 | 5902 | 994 | 1153 | 440 | 463 | 99 | 0 |
| Rain | 59181 | 16132 | 6890 | 1495 | 237 | 0 | 0 | 0 |
